# TRPM4 is required for calcium oscillations downstream from the stretch-activated TRPV4 channel

**DOI:** 10.1101/2021.12.15.472700

**Authors:** Oleg Yarishkin, Tam T. T. Phuong, Felix Vazquez-Chona, Jacques Bertrand, Sarah N. Redmon, Monika Lakk, Christopher N. Rudzitis, Jackson M. Baumann, Eun Mi Hwang, Darryl Overby, David Križaj

## Abstract

Transduction of mechanical information is influenced by physical, chemical and thermal cues but the molecular mechanisms through which transducer activation shapes temporal signaling remain underexplored. In the present study, electrophysiology, histochemistry and functional imaging were combined with gene silencing and heterologous expression to gain insight into calcium signaling downstream from TRPV4 (Transient Receptor Potential Vanilloid 4), a stretch-activated nonselective cation channel. We show that trabecular meshwork (TM) cells, which employ mechanotransduction to actively regulate intraocular pressure, respond to the TRPV4 agonist GSK1016790A with fluctuations in intracellular Ca^2+^ concentration ([Ca^2+^]_i_) and an increase in [Na^+^]_i_. ([Ca^2+^]_i_ oscillations coincided with a monovalent cation current that was suppressed by BAPTA, Ruthenium Red and 9-phenanthrol, an inhibitor of TRPM4 (Transient Receptor Potential Melastatin 4) channels. Accordingly, TM cells expressed TRPM4 mRNA, protein at the expected 130-150 kDa and showed punctate TRPM4 immunoreactivity at the membrane surface. Genetic silencing of TRPM4 antagonized TRPV4-evoked oscillatory signaling whereas TRPV4 and TRPM4 co-expression in HEK-293 cells reconstituted the oscillations. Membrane potential recordings indicated that TRPM4-dependent oscillations required release of Ca^2+^ from internal stores. 9-phenanthrol did not affect the outflow facility in mouse eyes. Collectively, our results show that TRPV4 activity initiates dynamic calcium signaling in TM cells by stimulating TRPM4 channels and intracellular Ca^2+^ release. These findings provide insight into the complexity of membrane-cytosolic interactions during TRPV4 signaling and may foster strategies to promote homeostatic regulation and counter pathological remodeling within the conventional outflow pathway of the mammalian eye.

## Introduction

Biomechanical factors such as IOP are an important determinant of the ocular environment, with roles in growth, cellular signaling that subserve ocular health and pathology (1)(2). IOP is homeostatically regulated by the trabecular meshwork (TM), a mechanosensitive phagocytic tissue composed of smooth muscle-, macrophage- and fibroblast-like cells that function as an adaptive valve for outflow of aqueous humor from the anterior eye (3)(4). In healthy cells, acute IOP increases engender an adaptive response that guides gradual recovery of tissue permeability to fluid flow (5) whereas chronic mechanical stress triggers transdifferentiation of TM cells into stiff and contractile myofibroblasts that subserve a lasting increase in outflow resistance (6)(7)(8). The appearance of myofibroblasts within the profibrogenic environment is often associated with inflammatory signaling and reduced aqueous humor outflow. Recent studies showed that TM cells sense acute and chronic mechanical stressors via arrays of mechanosensitive proteins that include integrins, the cytoskeleton, and stretch-activated ion channels (SACs) (9)(10)(11). Stretch-induced activation of TRPV4, a nonselective cation channel (P_Ca_/P_Na_ ~6-10), results in Ca^2+^ influx that regulates cytoskeletal and lipid dynamics, Rho signaling, cell-ECM interactions in TM cells and fluid outflow (12)(13)(14). Excessive TRPV4 activity was suggested to drive myofibroblast transdifferentiation that subserves the increase in TM stiffness and contractility in glaucoma (7)(14)(2) yet how channel signals in response to acute sustained activation remains unknown.

The human TRPV4 gene contains 16 exons that encode a protein consisting of a proline-rich (PRD) domain, phosphoinositide binding site, ankyrin repeats and a helix-turn-helix (HTH) linker domain within the N-terminus, six transmembrane domains with the pore between S5-S6, and modulatory sites (TRP domain, MAP7-binding domain, CaM/IP3R-binding domain, PDZ domain) in the C-terminus that endow the channel with sensitivity to temperature, metabolites of arachidonic acid, nociceptive and mechanical stimuli (15)(16). Gain- and loss-of-function mutations in the TRPV4 gene cause debilitating skeletal abnormalities, sensorimotor neuropathies and vision loss (17), indicating that the channel is required for homeostatic transduction of sensory information. TRPV4 overactivation and exposure to TGFβ induce transdifferentiation of epithelial, endothelial, and smooth muscle cells into hypersecretory and contractile myofibroblasts (18)(19). Interactions with a wide array of proteins (aquaporins, BK channels, kinases, actin) and processes (Ca^2+^ release from intracellular stores, receptor-operated and store-operated Ca^2+^ influx) (20)(21)(22)(13)(23) indicate that TRPV4 exerts its functions within a multiplicity of biological contexts.

The aim of this study was to simulate sustained application of mechanical stress by stimulating TRPV4 channels in the absence of confounding effects of the *in situ* milieu and parallel activation of TREK-1 and Piezo1 channels (10)(11). We report that continual chemical stimulation of TRPV4 induces [Ca^2+^]_i_ fluctuations that require TRPM4, a Ca^2+^-activated monovalent cation channel (P_Ca_/P_Na_ = 0.05) that has previously been linked to hypertension and Ca^2+^ oscillations (24)(25)(26)(27) and Ca^2+^ release from intracellular compartments. These findings suggest that TM mechanosensing involves dynamic interactions between membrane and cytosolic signaling mechanisms that endow the cells with the capacity for endogenous generation of time-dependent calcium signals.

## Materials and Methods

### Primary cell isolation and culture

Primary trabecular meshwork cells (pTM) were isolated from juxtacanalicular and corneoscleral regions of the human donors’ eyes (56 years-old male, 62 years-old female) with no recorded history of eye disease as described (13)(10)(11). The tissues were obtained through the Lions Eye Bank at the Moran Eye Center at the University of Utah and were used in concordance with the tenets of the WMA Declaration of Helsinki and the Department of Health and Human Services Belmont Report. A subset of biochemical experiments was conducted in parallel with immortalized cells (hTM), isolated from the juxtacanalicular region of the human eye (ScienCell Research Laboratories, Carlsbad, CA, USA). TM cells were cultured in accordance with consensus characterization recommendations, with phenotype periodically profiled for expression of *MYOC, TIMP3, AQP1, MGP, ACTA2* (α-smooth muscle actin, αSMA) genes and DEX-induced upregulation of myocilin protein (14)(11).

Passage 2 – 6 cells were seeded onto Collagen I-coated coverslips and grown in Trabecular Meshwork Cell Medium (ScienCell, Catalog#6591; Carlsbad, CA, USA) supplemented with 2% fetal bovine serum (FBS), 100 units/ml penicillin, 100 ug/ml streptomycin, at 37 ^o^C and pH 7.4. Cells were transfected with scrambled shRNA (Sc-shRNA) or *Trpm4* shRNA using Lipofectamin™ 3000 (5 μg per 25 cm^2^ tissue culture flask). shRNA - transfected cells were identified by green (GFP) or red (mCherry) fluorescence. Knockdown efficiency of TRPM4 shRNA was validated in HEK 293T cells overexpressing GFP-hTRPM4, scrambled shRNA or one of three TRPM4 shRNA constructs. shRNA1 and shRNA2, with target sequences GGACATTGCCCAGAGTGAACT (1218-1238) and GGAAAGACCTGGCGTTCAAGT (1835-1855), showed >65% knockdown efficiency. Experiments were conducted 3 - 4 days post-transfection.

### Reagents

The TRPV4 agonist GSK1016790A (GSK101) and antagonist HC067047 (HC-06) were purchased from Sigma (St. Louis, MO) or Cayman Chemical (Ann Arbor, MI). Salts were purchased from Sigma (St. Louis, MO) or VWR (Radnor, PA), CBA (4-chloro-2-[[2-(2-chlorophenoxy) acetyl aminobenzoic acid from Tocris Bioscience (Minneapolis, MN), and BAPTA (1,2-Bis(2-aminophenoxy)ethane-N,N,N’,N’-tetraacetic acid) from Alfa Aesar (Tewksbury, MA). GSK101 (1 mM), and HC-06 (20 mM) DMSO stocks were diluted in extracellular saline (98.5 mM NaCl, 5 mM KCl, 3 mM MgCl_2_, 2 mM CaCl_2_, 10 mM HEPES, 10 mM D-glucose, 93 mM mannitol), with final DMSO concentration not exceeding 0.1%.

### Immunocytochemistry

#### A. Cells

Cells were plated on collagen Type 1-coated glass cover slip 1 day prior to fixation with the 4% PFA for 10 minutes at RT. The samples were blocked with phosphate-buffered saline containing 0.03% Triton X-100 and 5% FBS for 30 minutes at a room temperature (RT). The primary polyclonal rabbit antibody was as described in Cho et al. (2014), diluted 1:1000 in PBS (2% BSA and 0.2% Triton X-100) and incubated overnight at 4^□^C. After rinsing, cells were incubated for 1 hour at RT with goat anti-mouse and goat anti-rabbit IgG (H + L) secondary antibodies conjugated to fluorophores (Alexa Fluor 488 nm, 568 nm and/or 594 nm; 1:500; Life Technologies, Carlsbad, CA, USA). Unbound antibody was rinsed, and conjugated fluorophores protected with Fluoromount-G (Southern Biotech, Birmingham, AL, USA) prior to mounting coverslips. Images were acquired on a confocal microscope (FV1200; Olympus, Center Valley, PA) at 1024 x 1024 pixels with a 20x water superobjective (1.00 N.A.; field size: 158.565 x 158.565 μm; 0.155 μm/pixel; sampling speed: 10.0 us/pixel; 12 bits/pixel). ≥50 cells per experiment were acquired for at least 4 independent experiments (n ≥ 200 in total; N ≥ 4).

#### B. Tissue immunohistochemistry

Anterior chambers were fixed in 4% paraformaldehyde for 1 hour, cryoprotected in 15 and 30% sucrose gradients, embedded in Tissue-Tek^®^ O.C.T. (Sakura, 4583), and cryosectioned at 12 μm, as described (13)(27). Sections were probed with a polyclonal rabbit TRPM4 antibody (1:100; (28) and AQP1 mouse monoclonal antibody (1:1000; Santa Cruz Biotechnology Sc-25287). Secondary antibodies were anti-rabbit IgG DyLight 488 (Invitrogen, 35552) and anti-mouse IgG DyLight 594 (Invitrogen, 35511). Sections were coverslipped with DAPI-Fluoromount-G (Electron Microscopy Sciences, Hatfield, PA, 17984-24) and imaged with a confocal microscope. Images were acquired using identical (HV, gain, offset) parameters.

### Semiquantitative Real Time-PCR

Total RNA was extracted using Arcturus PicoPure RNA Isolation Kit (ThermoFisher, Cat. No.: KIT0204) (29)(30). 100 ng of total RNA was used for reverse transcription. First-strand cDNA synthesis and PCR amplification of cDNA were performed using qScript™ XLT cDNA Supermix (Quanta Biosciences, Cat. No.: 95161). The samples were run on 2% agarose gels using ethidium bromide staining along with the 100-bp DNA ladder (ThermoFisher Scientific, Waltham, MA, Cat. No.: S0323). The primers used in the study are listed in Table I.

### Western blot

TM cells were detached from culture flasks by trypsinization and centrifuged at 2000 rpm for 3 minutes. The cell pellet was washed with PBS and lysed in a RIPA Buffer System (Santa Cruz Biotechnology, Dallas, TX). Cell lysates were separated by 10% SDS-PAGE followed by electrophoretic transfer to polyvinylidene difluoride membranes (Bio-Rad, Hercules CA). Membranes were blocked with 5% skim milk in PBS containing 0.1% Tween 20 and incubated at 4°C overnight with the TRPM4 antibody. After developing with TRPM4 antibody, membranes were washed 3 times with PBS containing 0.1% Tween-20, blotted with HRP-conjugated GAPDH antibody for 30 min at room temperature and washed with PBS containing 0.1% Tween 20. The signals were visualized with an enhanced chemiluminescence system (FluorChem Q, Cell Biosciences, Santa Clara, CA).

### Ion imaging

Pharmacological experiments were conducted on a microscope stage in a fast-flow chamber (RC26GLP, Warner Instruments, Hamden CT) connected to a gravity-fed perfusion system. The flow rate of the solution was regulated via individual pinch valves (VC-6; Warner Instruments; Holliston, MA). Primary cells were loaded with Fura-2 AM (3 μM; Invitrogen/ThermoFisher Scientific) for 45 min at RT. The cells were perfused with extracellular solution containing (mM): 135 NaCl, 2.5 KCl, 1.5 MgCl_2_, 1.8 CaCl_2_, 10 HEPES, 5.6 D-glucose (pH = 7.4, osmolarity = 300 - 303 mOsm). Epifluorescence imaging was performed on inverted Nikon microscopes using 40x (1.3 N.A., oil or 0.80 N.A., water) objectives, 340 nm and 380 nm excitation filters (Semrock, Lake Forest, IL, USA) and a Xenon arc lamp (DG4, Sutter Instruments). Fluorescence emission at 510 nm, in response alternate 340/340 excitation, was captured with cooled EMCCD or CMOS cameras (Photometrics, Tucson, AZ). Data acquisition was controlled by NIS Elements 3.22 software (Nikon). Typically, ~5-10 cells per slide were averaged across ~3-6 slides per experiments, with at least 3 independent experiments (n ≥ 50; N ≥ 3). The experiments were performed at room temperature (20 - 22° C).

#### Ca^2+^ imaging

Cells were loaded with 3 μM Fura-2 AM (Invitrogen/ThermoFisher) (K_d_ at RT = 225 nM) for 45 min. ΔR/R (peak F_340_/F_380_ ratio – baseline/baseline) was used to quantify the amplitude of Ca^2+^ signals (e.g., (13)(31). Only transient Ca^2+^ events with amplitudes exceeding three standard deviations of baseline fluctuations were included in analysis.

#### Intracellular Na^+^ imaging

Cells were loaded with 3 μM NaTRIUM Green™-2 AM (TEFLabs Austin, TX) for 50 – 60 min in a cell culture incubator at 37^0^C. Fluorescence was acquired at 484 nm excitation and 520 nm emission (Semrock, Rochester, NY). Na^+^ signals were normalized to the average baseline (F/F_o_) obtained at the beginning of the experiments.

### Electrophysiology

Whole cell and single-channel techniques were used to record membrane currents (voltage clamp) or voltage (current clamp) (10)(32)(11). Borosilicate patch pipettes (WPI, Sarasota, FL) were pulled to resistances of 5 - 8 MΩ (P-2000; Sutter Instruments, Novato CA). The standard pipette solution contained (in mM): 125 NaCl, 10 KCl, 10 HEPES, 1 MgCl_2_, 2 ethylene glycol-bis(β-aminoethyl ether)-N,N,N’,N”-tetraacetic acid (EGTA), 0.3 Na-GTP (pH 7.3) The pipette solution for recording Ca^2+^-activated current contained (in mM): 125 Na-gluconate, 10 HEPES, 1 MgCl_2_, 0.01 CaCl_2_, 0.3 Na-GTP (pH 7.3). Nominally Ca^2+^-free pipette solution (standard) contained (in mM): 125 Na-gluconate, 10 HEPES, 1 MgCl_2_, 5 BAPTA, 0.3 Na-GTP (pH 7.3). The standard extracellular solution contained (in mM): 135 NaCl, 2.5 KCl, 1.5 MgCl_2_, 1.8 CaCl_2_, 10 HEPES, 5 D-glucose (pH 7.4). The extracellular solution for recording Ca^2+^-activated current contained (in mM): 135 Na-gluconate, 10 HEPES, 1 MgCl_2_, 1.8 CaCl_2_, 0.3 Na-GTP (pH 7.3). The bathing solution for excised (inside-out) patch clamp experiments contained (mM): 140 NaCl, 2.5 KCl, 1.5 MgCl_2_, 1.8 Ca^2+^ or 10 EGTA, 5.6 D-glucose, 10 HEPES (pH 7.4, adjusted with NaOH). The pipette solution used in excised patch recordings contained 140 NaCl, 2.5 KCl, 1.5 MgCl_2_, 1.8 CaCl_2_, 5.6 D-glucose, 10 HEPES (pH 7.4, adjusted with NaOH). Experiments were performed at room temperature of 21-22 °C.

Patch clamp data in whole-cell and single-channel configurations was acquired with a Multiclamp 700B amplifier and a Digidata 1550 interface, and controlled by Clampex 10.7 (Molecular Devices, Union City, CA). Cells were held at −100 mV or +100 mV, with currents sampled at 5 kHz, filtered at 2 kHz with an 8-pole Bessel filter and analyzed with Clampfit 10.7 (Molecular Devices) and Origin 8 Pro (Origin Lab, Northampton, MA). Whole-cell currents were elicited by voltage ramps ascending from −100 mV to 100 mV (0.2 V/sec) from the holding potential of 0 mV. The time course of Ca^2+^-activated currents was obtained by subtracting voltage ramp-evoked currents traces recorded 40 s after obtaining the whole-cell configuration from all preceding and following traces. I-V curves of Ca^2+^-activated currents were assessed by subtracting currents recorded 40 s after obtaining the whole-cell configuration from steady state currents recorded 4 min after obtaining the whole-cell configuration.

### Patch clamp combined with fluorescent imaging

Cells were loaded with a calcium indicator Fluo-4 (Invitrogen/ThermoFisher) through a patch pipette. Fluorescence signals in voltage-clamped cells were detected using the excitation filter 484 nm and the emission filter 520 nm (both from Semrock, USA). VOCC-mediated current was recorded in the extracellular solution containing as following (mM): 115 NaCl, 2.5 KCl, 20 BaCl_2_, 1.5 MgCl_2_, 10 HEPES, 5.6 D-glucose.

### Outflow facility measurement

Procedures on living mice were carried out under the authority of a UK Home Office project license and adhered to the ARVO Statement for the Use of Animals in Ophthalmic and Vision Research.

### iPerfusion

The outflow facility in enucleated eyes from C57BL/6J mice (N = 5 mice; 11-week-old males; Charles River UK Ltd., Margate, UK) was measured using *iPerfusion*, a custom-made microfluidic setup (33)(34). Following cervical dislocation, eyes were enucleated and affixed to a support platform using tissue glue. Eyes were submerged in a PBS bath kept at 35°C throughout the perfusion. Using a micro-manipulator under a dissection microscope, the anterior chamber was cannulated within 10 minutes of death using a glass micropipette pulled to have a 100 μm diameter beveled tip. The perfusion fluid was Dulbecco’s PBS containing divalent cations and 5.5 mM glucose (DBG) that was sterile filtered (0.25 μm) prior to use.

One eye was perfused with 9-PA (25 μM) whilst the contralateral eye was perfused with vehicle (perfusion fluid containing the same concentration of DMSO). IOP was set to 9 mmHg for 1 hour to pressurize and acclimatize the eye to the perfusion environment and to allow sufficient time for 9-PA to reach the outflow tissues. Flow into the eye was then measured over 8 increasing pressure steps from 6.5 to 17 mmHg. Steady state for each step was evaluated when the ratio of the flow rate to pressure changed by less than 0.1 nl/min/mmHg per minute over a 5-minute window (33). The stable pressure, ***P***, and flow rate, ***Q***, were calculated over the last 4 minutes of each step, and a power-law relationship of the form

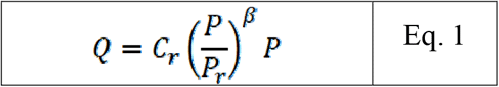

was fit to the ***Q – P*** data. The reference outflow facility, ***C_r_***, represents the value of outflow facility at a reference pressure ***P_r_*** of 8 mmHg, and ***β*** characterizes the non-linearity of the ***Q – P*** relationship (33). The relative difference in ***C_r_*** between treated and untreated contralateral eyes was then calculated as the ratio of ***C_r_*** in the treated eye relative to that in the contralateral control eye minus unity. We evaluated whether the relative difference in facility was statistically different from zero using a weighted *t*-test on the log-transformed data, as previously described (33). Facility values and relative changes in facility are reported in terms of the geometric mean and the 95% confidence interval on the mean.

### IOP measurement

A TonoLab rebound tonometer was used to measure IOP of Avertin-treated mice between noon and 2 PM. An IOP reading represents an average of 10 to 20 tonometer readings.

### Statistical Analysis

Student’s paired t-test or two-sample t-test were applied to estimate statistical significance of results. P < 0.05 was considered statistically significant. Results are presented as the means ± S.E.M.

## Results

### Activation of TRPV4 induces fluctuations of [Ca^2+^]_i_

Unstimulated TM cells from healthy donors have low basal [Ca^2+^]_i_ and rarely show Ca^2+^ fluctuations (13) but little is known about Ca^2+^ homeostasis in the presence of sustained TRPV4 activation. Previous studies of mechanically evoked Ca^2+^ signaling in TM cells were limited to short stimulation paradigms (5-10 min) (13)(11). To investigate the mechanisms that underlie continual Ca^2+^ signaling downstream from the putative mechanotransducer TRPV4, Fura-2-loaded human TM cells were stimulated for 30 – 45 min with bath-applied agonist GSK101 (25 nM). [Ca^2+^]_i_ increases evoked by the agonist peaked within 2-3 min and were followed by gradual decline to a steady plateau at ~27% of the peak (Figs. 1 & 2). A subset (~50%) of cells showed an increase in the frequency of Ca^2+^ fluctuations during the plateau phase whereas cells treated with the TRPV4 antagonist HC067047 (HC-06; 5 μM) (Fig. 1B & C), broad-spectrum inhibitors of TRP channels such as RuR, or Ca^2+^ -free saline (Fig. 2), did not exhibit this timedependent behavior.

**Figure 1.**
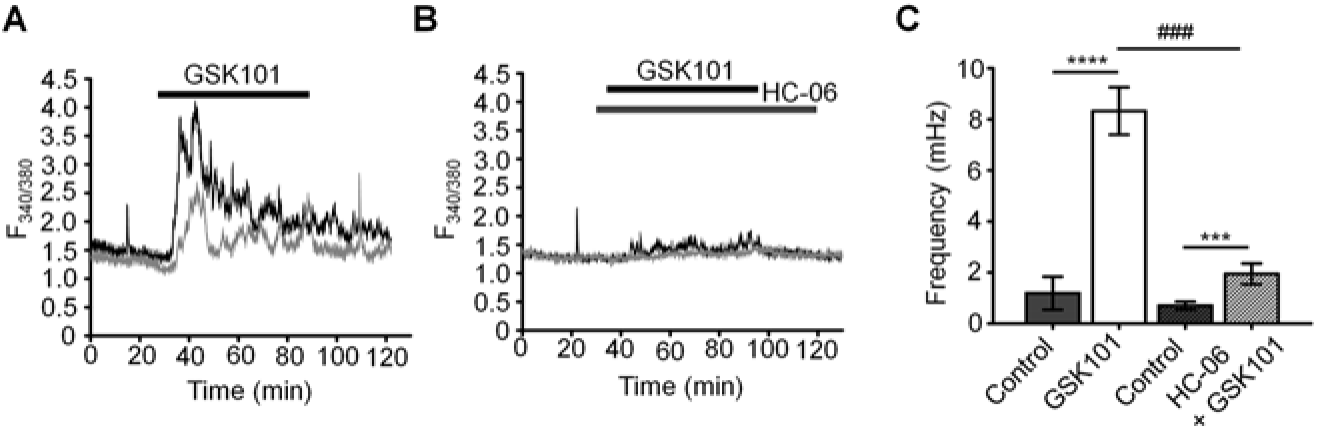
GSK101 triggers intracellular Ca^2+^ oscillations in hTM cells. **(A)** A representative trace of [Ca^2+^]_i_ oscillations in GSK101 treated TM cells. **(B)** GSK101-induced oscillations are absent in cells pretreated with HC-06. **(C)** Quantification of results shown in (A) and (B), shown as average ± S.E.M. ****P < 0.0001,***P < 0.001 (paired-sample t-test), ###P < 0.0001 (two-sample t-test), n = 49 cells and n = 38 cells for control/GSK101 and control/HC06+GSK101 groups, respectively.

**Figure 2.**
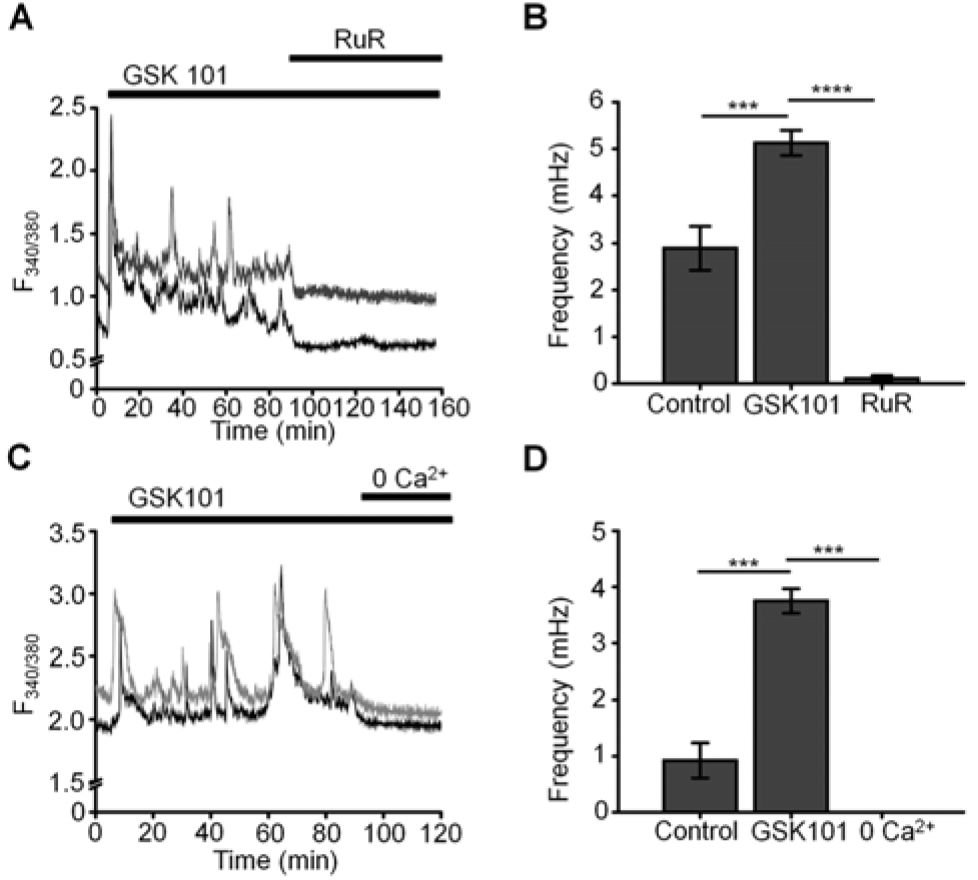
GSK101-induced [Ca^2+^]_i_ oscillations requires activity of Ca^2+^ -permeable and Ca^2+^ -dependent TRP channels. **(A and C)** GSK101-induced [Ca^2+^]_i_ fluctuations are abolished by the nonselective TRP blocker Ruthenium Red (RuR; 10 μM) and in the absence of free extracellular Ca^2+^. **(B and D)**RuR and Ca^2+^-free solution abolish GSK101 induced [Ca^2+^]_i_ oscillations. Mean ± S.E.M. ***P < 0.001, ****P < 0.0001, paired t-test, n = 51 and n = 43 cells for (B) and (D), respectively.

Oscillatory Ca^2+^ signals can reflect activation of voltage-operated mechanisms within the membrane and/or release from internal compartments. Voltage-dependent mechanisms were tested by stimulating the cells in the current-clamp mode. The agonist evoked a ~15 mV shift in the membrane potential (Fig. 3A & C) that was maintained over the course of the experiment (e.g., 40 min after agonist application; red trace in Fig. 3B) and was not associated with changes in temporal behavior.

**Figure 3.**
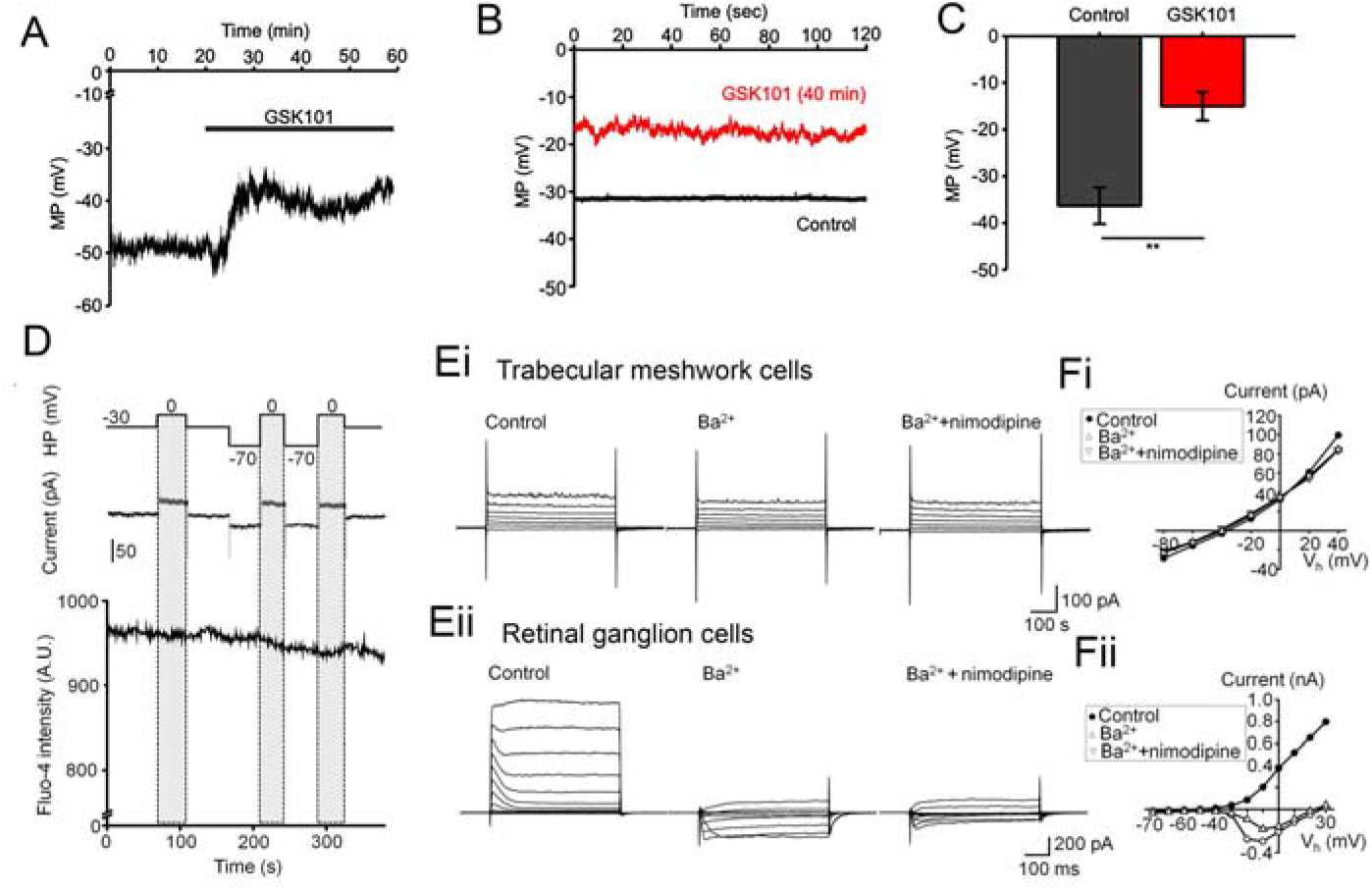
GSK101 depolarizes TM cells, but they lack functional L-type Ca^2+^ channels. **(A - C)** Current clamp. GSK101 depolarizes the cells, with the effect maintained over 40 min (red trace vs. black trace denoting the unstimulated control in B). **(D)** Combined whole-cell recording and calcium imaging. The voltage protocol (upper trace), depolarization-evoked whole-cell current (middle trace), time-lapse of Fluo-4 fluorescence (lower trace). Shadowed bars indicate the depolarizing steps. **(E & F)** Representative whole-cell currents elicited by voltage steps in TM cells (Ei; n = 15) (B) and mouse retinal ganglion cells (Eii; n = 5), respectively. **(F)** Voltage-current relationships derived from cells shown in (E). The current was measured at the end of each voltage pulse.

L-type voltage-gated Ca^2+^ channels (VGCCs) have been implicated in calcium oscillations and pressure-dependent contractility in smooth muscle cells (35). To test their involvement, we combined imaging in Fluo-4-loaded cells with whole-cell recording. As shown in Figure 3D, stepping the V_m_ from the holding potential of −70 mV to 0 mV had no detectable effect on the Ca^2+^ signal. Similarly, extracellular BaCl_2_ (20 mM) which permeates voltage-operated Ca^2+^ channels (VOCCs) without inactivating the channel pore or stimulating Ca^2+^ release from ER stores (36)(37)(38) did not induce depolarization-evoked inward currents or Ca^2+^ spiking (Fig. 3Ei) whereas retinal ganglion cells responded with an inward current (Fig. 3E, F) that was blocked by the L-type channel blocker nimodipine (10 μM) (Fig. 3Fii). These data suggest that TRPV4-dependent Ca^2+^ oscillations in human TM cells do not involve VGCC activity.

### TRPV4-induced Ca^2+^ fluctuations require TRPM4

To further investigate the mechanism that underlie TRPV4-induced calcium oscillations, cells were loaded with the Na^+^-sensitive indicator NaTRIUM Green-2 and exposed to GSK101. The agonist stimulated an increase in [Na^+^]_i_ in 54% (39/73) cells (5.96 ± 0.81 % above baseline; p < 0.0001; n = 39). The increase in Na^+^ concentration was not associated with an oscillatory component, indicating that mechanisms that mediate TRPV4-induced Na^+^ influx are distinct from mechanisms that subserve Ca^2+^ oscillations. TRPM4, a Ca^2+^-activated channel permeable to monovalent ions, has been implicated in [Ca^2+^]_i_ oscillations in T cells, cardiomyocytes, and smooth muscle cells (35)(24)(25) with no known functions in the eye. To test its involvement in TRPV4-induced Ca^2+^ oscillations, we exposed the cells to GSK101 in the presence of 9-phenanthrol (9-PA, 40 μM), an intracellularly acting benzoquinolizinium inhibitor of the channel (39)(40). 9-PA reversibly attenuated the increase in [Na^+^]_i_ evoked by GSK101 (P < 0.001) (Fig. 4A & B), and obliterated Ca^2+^ fluctuations during the plateau response phase (Fig. 4 C&D) (P < 0.0001).

**Figure 4.**
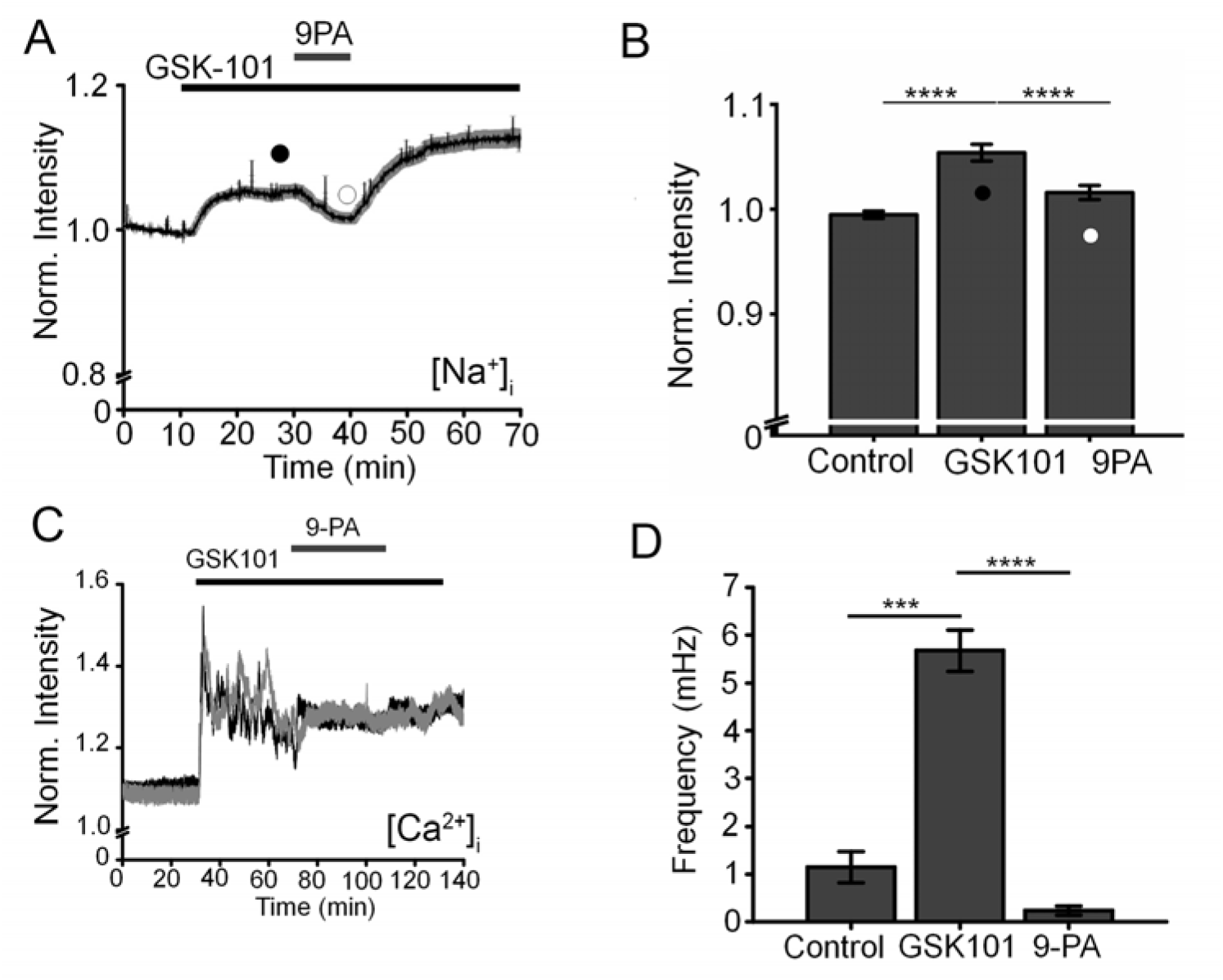
TRPM4 mediates TRPV4-induced Na^+^ influx and facilitates Ca^2+^ fluctuations. **(A)** Averaged time lapse ratio in NaTRIUM Green-loaded TM cells. GSK101-evoked increase in [Na^+^]_i_ is partially and reversibly suppressed by 9-PA. **(B)** Mean ± S.E.M. values for the experiment in A (n = 10). Black and white symbols represent time at which the intensity was measured. **(C**) GSK101-evoked [Ca^2+^]_i_ oscillations are suppressed by 9-PA (n = 39). **(D)** Summary of the data shown in C, as mean ± S.E.M. ** = p < 0.01, **** = p < 0.0001, paired-sample t-test.

### Histology: TM cells express TRPM4

Transcriptional profiling of immortalized (hTM) and primary (pTM) cells showed robust expression of TRPM4 and TRPV4 transcripts (Fig. 5A) together with TM markers MYOC and AQP1, at bp sizes that matched expected sizes for their respective DNAs. The overall levels of TRPM4 transcripts in hTM cells were comparable to TRPV4 mRNA, and ~35% of TRPV4 mRNA in primary cells (Fig. 5B). In both cell lines, a validated antibody (28) labeled ~130 kDa Western blot band (Fig. 5C) that corresponds to the known M.W. of the TRPM4b protein variant (41). Confocal examination of TRPV4-immunolabeled cells and intact tissue further showed labeling of the plasma membrane (inset of Fig. 5Dii) and cytosolic puncta. In the preparation from the mouse anterior eye, TRPM4-ir was distributed across the juxtacanalicular and corneoscleral TM regions (green), where it colocalized with the TM marker Collagen IV (Col IV; blue). Additional, albeit more modest, TRPM4-ir signals were also detected in the ciliary body and the retina.

**Figure 5.**
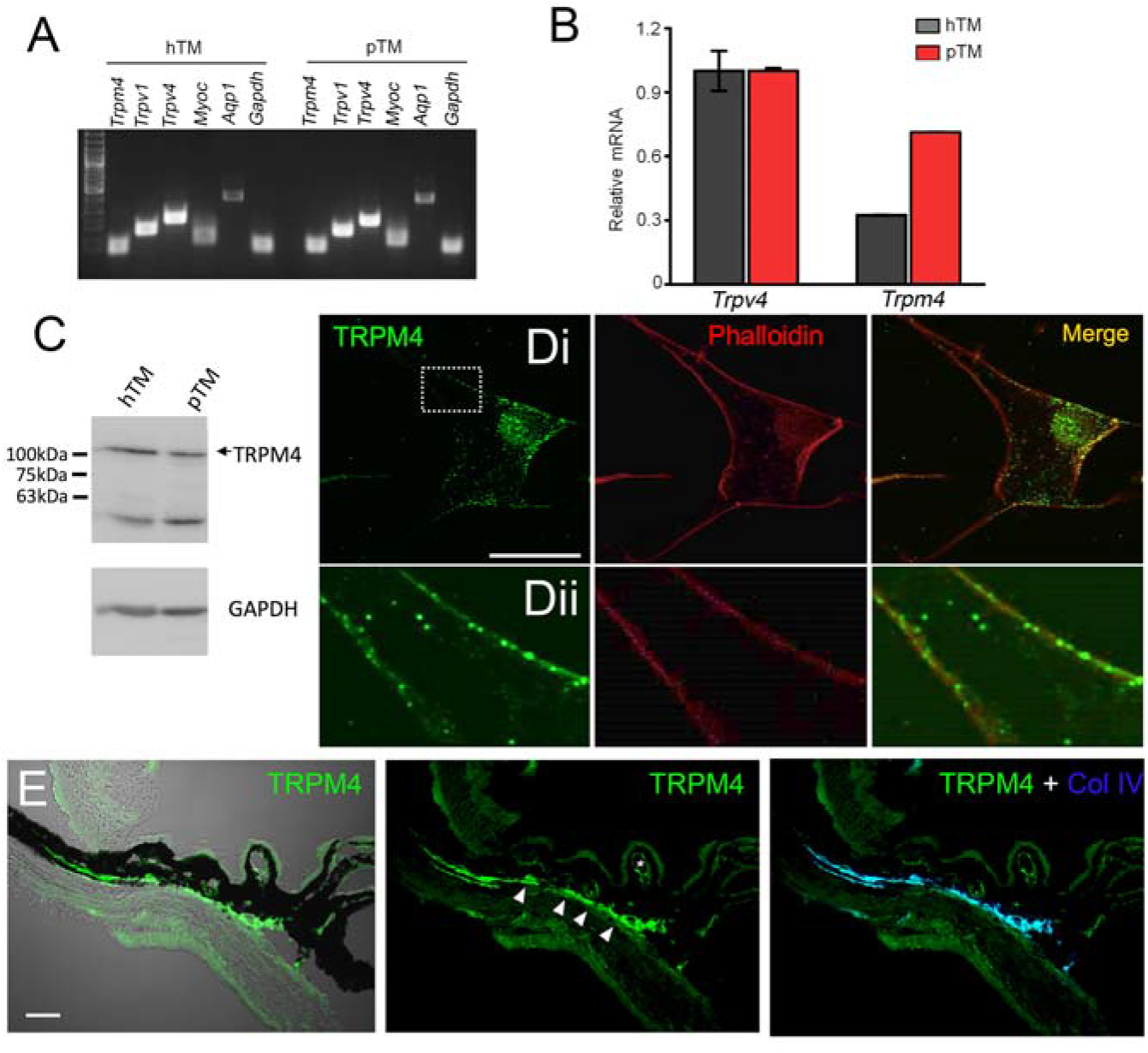
TRPM4 is expressed in TM cells. **(A)** Representative PCR gels show expression of TRPM4 and TRPV4 transcripts together with markers AQP1, TRPV1, and MYOC in immortalized (hTM) and primary (pTM) cells. **(B)** Semiquantitative qRT-PCR. Relative mRNA abundance of TRPM4 vs. TRPV4; expression is normalized to hTM TRPM4; mean ± S.E.M, N = 3. **(C)** Western blot. hTM and pTM cells show a TRPM4 band at the expected M.W. **(D,** *upper panel*) Cultured cells immunolabeled for TRPM4, with a magnified image of the region selected by the square (*inset*) showing punctate signals within the plasma membrane. Scale bar = 50 μm. **(E)** Anterior eye section from a wild type C57 mice, labeled with TRPM4. The TM region shows pronounced TRPM4-ir (arrows) that colocalizes with the marker Collagen IV. Moderate signal is seen in the ciliary body (arrows) and ciliary vasculature (asterisk).

### Electrophysiology: TM cells functionally express TRPM4

We next assessed the properties of the TRPM4-mediated current in voltage-clamped cells by probing for a Ca^2+-^ sensitive monovalent conductance (42) (43). Cells were dialyzed with pipette solutions containing 10 μM or zero Ca^2+^ and Ca^2+^-activated K^+^ and Cl^-^ conductances were minimized by replacing extra/intracellular K^+^ ions (with Na^+^) and Cl^-^ ions with gluconate (42). ~50% of cells dialyzed with the Ca^2+^-containing solution showed time-dependent increases in inward and outward whole-cell currents. The current-voltage (I-V) relationship of the Ca^2+^-dependent current showed modest outward rectification and peaked within ~4 min after obtaining the whole-cell configuration (Fig. 6A), had a reversal potential of 1.6 ± 2.9 mV and amplitudes of −122.7 ± 61.5 pA and 211.2 ± 93.7 pA at holding potentials of −100 mV and 100 mV, respectively, (Fig. 6B). The current was inhibited by 9-PA (20 μM; n=7/7) and was not observed in cells dialyzed with Ca^2+^-free solutions or exposed to bath-applied Ruthenium Red (RuR, 10 μM) (n = 5/5) (Fig. 6A).

**Figure 6.**
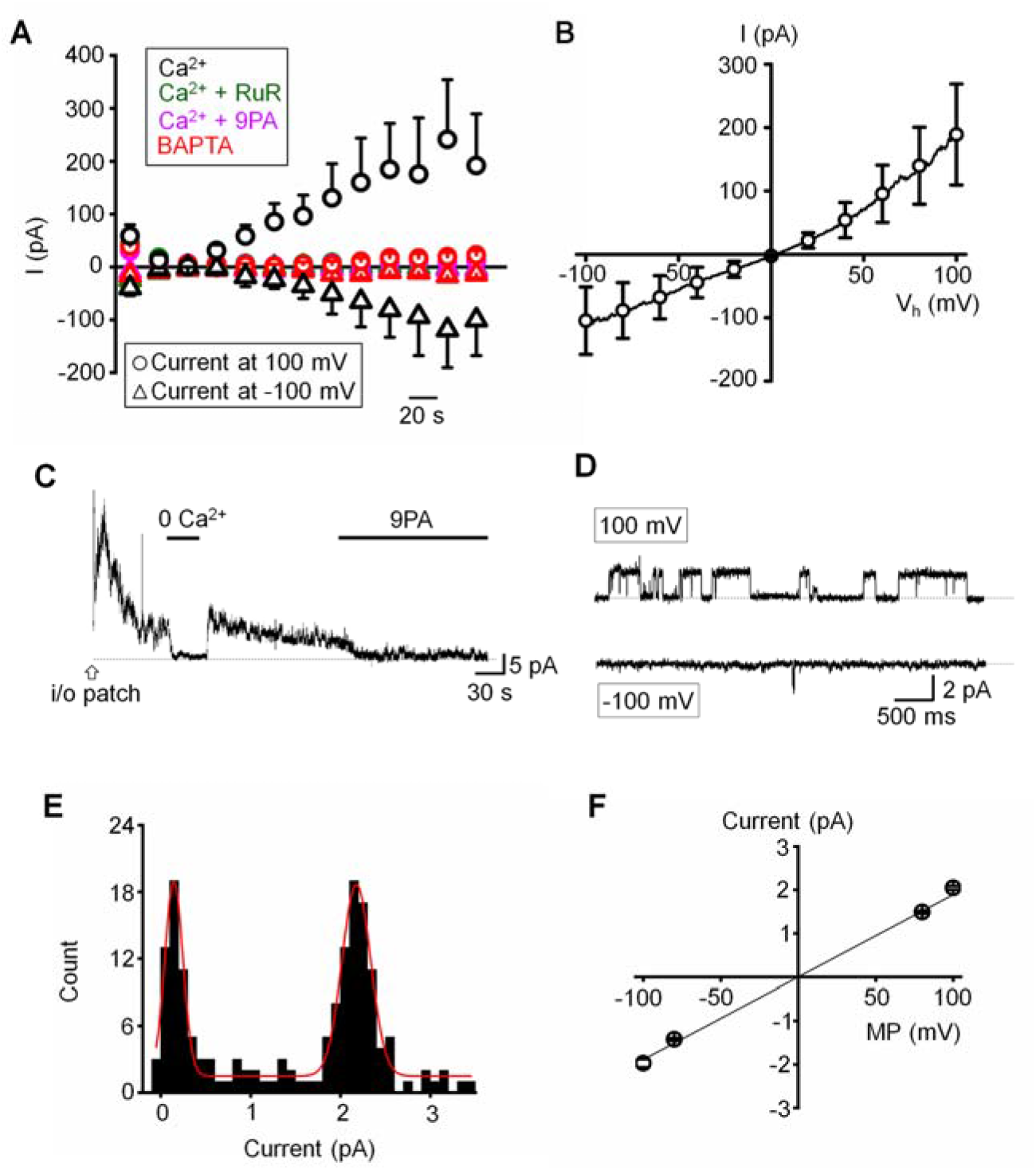
Functional expression of a calcium-activated CAN. **(A)** Time course of the whole-cell CAN. The pipette solution contained 10 μM Ca^2+^ in the standard saline (*black*, n = 5), with addition of RuR (10 μM; *olive*, n = 11), 9-PA (20 μM; *magenta*, n = 15) in extracellular solution, or Ca^2+^-free pipette solution (*red*, n = 15). **(B)** Averaged I-V relationship of the Ca^2+^ activated current in A subtracted by the Ca^2+^-free component (n = 6). **(C)** Representative trace demonstrating the activity of a Ca^2+^-activated channel in an inside-out patch preparation at +100 mV. **(D)** Representative traces of single channel activity. **(E)** Histogram of unitary current amplitude, fit with Gaussian function (red trace; R-Square = 96.6%). The data were binned at 0.02 pA. **(F)** Current-voltage relationship of single channel current, n = 6 patches. (A), (B) and (F) show mean ± S.E.M.

Single channel properties of the Ca^2+^-activated current were additionally investigated in inside-out membrane patches using symmetrical Na^+^ gluconate-based pipette and extracellular solutions. Patch excision in 1.8 mM Ca^2+^ -containing saline triggered a rapidly inactivating channel with a linear I-V relationship (n = 15/85 patches; Fig. 6C&F). The slope conductance of the unitary current was 18.9 ± 0.6 pS and the amplitude at 100 mV was 2.05 ± 0.04 pA (Fig. 6D-F). The channel was facilitated at positive membrane potentials, inhibited by 9-PA (Fig. 6C&D), and typically inactivated within ~5 min after patch excision. Its activity was abolished in Ca^2+^-free saline (Fig. 6C). The whole-cell and single channel properties of the Ca^2+^-induced current in TM cells are thus consistent with TRPM4.

#### TRPM4 is required for TRPV4-dependent Ca2+ oscillations

To more specifically test the role of TRPM4 in [Ca^2+^]_i_ fluctuations, we transfected the cells with TRPM4-specific short-hairpin (shTRPM4-GFP) or “scrambled” (Sc-GFP) control shRNAs (Fig. 7A). Of the 3 constructs, we used construct Sh#1 which downregulated TRPM4 mRNA by 80%. Compared to Sc-shRNA-expressing controls, TRPM4 shRNA-transfected cells showed ~80% reduction in the frequency of TRPV4-induced Ca^2+^ fluctuations (Fig. 7B-D).

**Figure 7.**
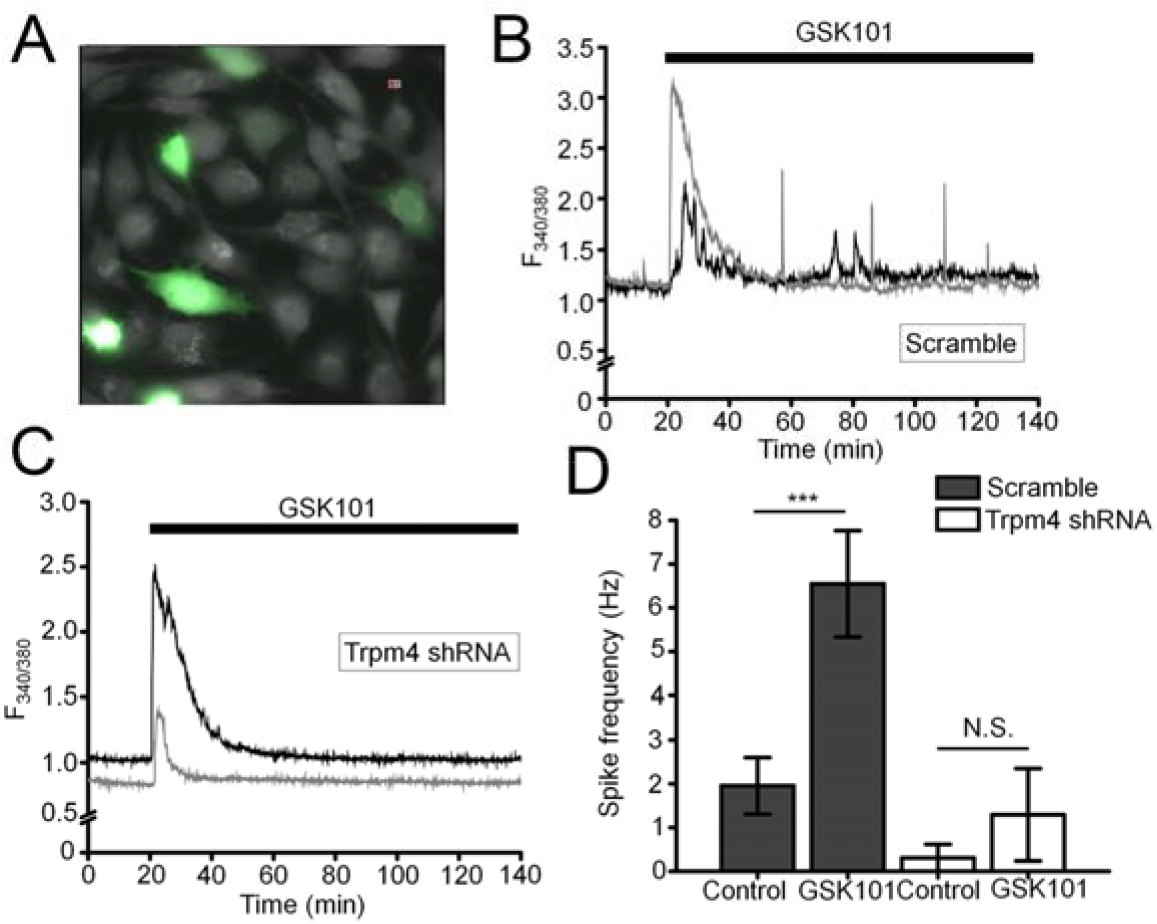
TRPM4 knockdown disrupts GSK101-triggered [Ca^2+^]_i_ fluctuations. **(A)** TM cells treated with Sc-GFP, and TRPM4 shRNA-GFP constructs. **(B & D)** Sc-shRNA-transfected cells exhibit fluctuating [Ca^2+^]_i_ signals, which show **(C & D)** reduced frequency in TRPM4-shRNA-treated cells. Data shown as mean ± S.E.M. ***P < 0.001, ^N.S.^P > 0.05, paired t-test, n = 116 cells and n = 27 cells for Sc- and TRPM4 shRNAs, respectively.

We additionally tested the TRPM4-dependence of TRPV4-induced calcium signals by transfecting HEK293 cells with TRPV4 and/or TRPM4 DNA. TRPV4 only-transfected cells responded to GSK101 with a peak [Ca^2+^]_i_ increase that inactivated to a plateau without showing time-dependent Ca^2+^ oscillations whereas cells co-transfected with TRPV4 + TRPM4 DNA displayed robust oscillations in the presence of GSK101.

#### TRPM4 does not regulate trabecular outflow or steady-state IOP

Chronic TRPV4 activation was suggested to suppress aqueous humor drainage in hypertensive mouse eyes (13). Given that 9-PA inhibits TRPV4-dependent Ca^2+^ oscillations in vitro (Fig. 8A & B), we tested whether it influences the trabecular component of pressure-induced fluid outflow (“outflow facility”). The effect of 9-PA on outflow facility was examined in enucleated mouse eyes using the *iPerfusion* system (33)(34). In response to 25 μM 9-PA, outflow facility decreased by −5% [-23%, 18%] (geometric mean [95% CI]; Fig. 8A) relative to contralateral eyes that were perfused with vehicle, but this difference was not statistically significant (*p*=0.38, n=5 pairs). ***C_r_*** for 9-PA treated eyes was 5.4 [4.2, 6.9] nl/min/mmHg and 5.5 [4.6, 6.7] nl/min/mmHg for vehicle-treated eyes (Fig. 8A).

**Figure 8.**
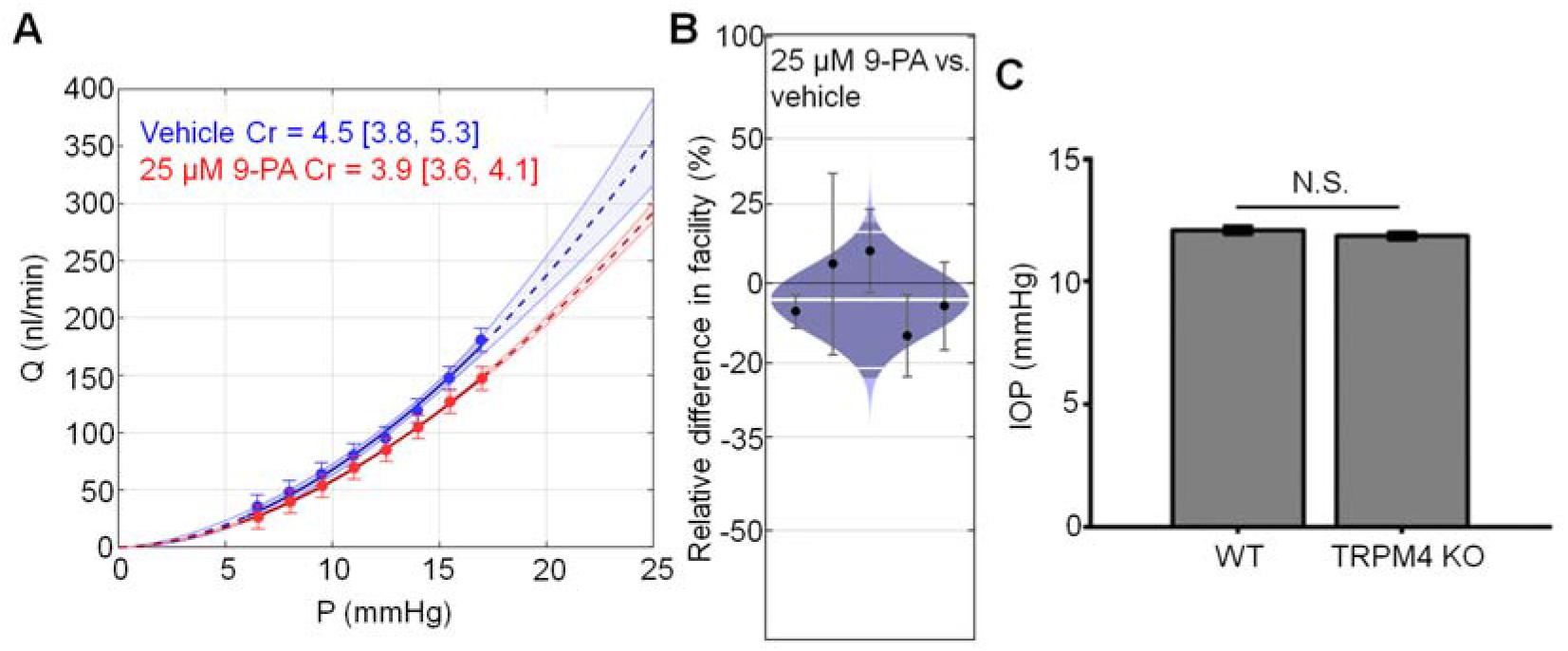
TRPM4 does not regulate trabecular outflow and steady-state IOP. **(A)** Representative flow-pressure ***(Q – P)*** plot for contralateral eyes perfused with 25 μM 9-PA versus vehicle. Curves show the optimal fit from Equation 1, with 95% confidence bounds and error bars show 95% confidence intervals. **(B)** Relative difference in outflow facility between contralateral eyes perfused with 25 μM 9-PA versus vehicle. The relative difference in facility is defined as the ratio of ***C_r_*** in the experimental eye perfused with 9-PA relative to vehicle-perfused contralateral eye minus unity, expressed as a percentage. Each data point represents the relative difference in facility for an individual mouse. Error bars are 95% confidence intervals. Shaded regions represent the best estimate of the sample distributions, with the central white line representing the geometric mean. Dark central bands represent the 95% CI on the mean, and the outer white lines represent the limits encompassing 95% of the population. **(C)** Summarized results for IOP in wild-type (WT, n = 14 eyes) and *TRPM4* KO (n = 36 eyes) mice. ^N.S.^ = P > 0.05.

We also tested whether genetic deletion of TRPM4 channels affects IOP homeostasis in the mouse eye. TRPM4^-/-^ eyes showed IOP levels (11.86 ± 0.14 mm Hg that were comparable to controls (12.10 ± 0.16 mm Hg; p > 0.05; n = 14 eyes and n = 36 eyes for WT and TRPM4^-/-^, respectively), indicating that TRPM4 may not be required for steady-state IOP regulation **(Fig. 8C)**. These results suggest that TRPM4 signaling is not required for step-induced fluid drainage across the or for steady-state IOP regulation.

## Discussion

The goal of this study was to define the properties of calcium homeostasis during sustained stimulation of TRPV4, a transducer of mechanical stimuli that has been implicated in the regulation of conventional outflow and IOP (13). We show that long-term TRPV4 stimulation evokes Ca^2+^ oscillations that are independent of the membrane potential and require expression of TRPM4. Both TRPV4 and TRPM4 were strongly expressed in primary and immortalized TM cells, with functional coupling between the two cation channels revealed by the Ca^2+^-activated monovalent current and loss of oscillatory response following TRPM4 knockdown. Our findings identify novel interactions between TRPV4, TRPM4 and intracellular calcium stores and build a new working model that may help increase our understanding of homeostatic and time-dependent signaling within the primary outflow pathway in the eye.

Changes in [Ca^2+^]_i_ represent one of the earliest responses of nonexcitable cells to mechanical stress. Resting human TM cells do not exhibit spontaneous calcium activity whereas sustained stimulation of TRPV4 channels produced three types of time-dependent calcium behavior. (i) The initial increase in [Ca^2+^]_i_ evoked by GSK101 peaked within 2-4 min and showed slow onset kinetics that was likely shaped by the messenger pathway involving obligatory activation of phospholipase A2 and production of epoxyeicosatrienoic messengers that bind the pocket formed by S2-S3/S4-S5 linker residues (13)(16). (ii) The subsequent relaxation to the steady-state plateau phase reflects Ca^2+-^dependent channel inactivation, internalization, interactions with modulatory sites within N- and C-termini (15) or negative regulation of Ca^2+^ influx by TRPM4 (43). (iii) The relaxation phase was associated with Ca^2+^ fluctuations in a frequency range similar to signals reported in glaucomatous TM (44) and pressure-, stretch- and GSK101-stimulated fibroblasts and chondrocytes (45)(46).

Several pieces of evidence suggest that the oscillatory Ca^2+^ response requires TRPM4 signaling. (i) TRPV4-induced fluctuations were suppressed by 9-PA, which also induced a reversible decrease in [Na^+^]_i_; (ii) TRPM4 knockdown in TM cells preserved the GSK101 response but eliminated the oscillatory calcium component, and (iii) Heterologous expression of TRPV4 and TRPM4 channels reconstituted fluctuations in HEK293 cells. TRPM4 expression in TM cells was confirmed by transcriptional profiling and immunohistochemistry, with *TRPM4* mRNA levels reaching ~30-70% of *TRPV4* mRNA, and immunoblots showing a band at the predicted M.W. As in other cell types (47)(48), the TRPM4 antibody labeled plasma membrane and intracellular compartments. The former corresponds to full-length TRPM4b variant which forms the Ca^2+^-activated Ca^2+^-impermeant cation channel and appears to congregate into punctate clusters within the cell membrane (Fig. 5Dii) whereas intracellular puncta may correspond to truncated N-terminal variants with unknown functions that localize to the ER (49)(50)(28). The aggregation of TRPM4-ir signals resembles mechanosensory clusters enriched in TRPV4, paxillin and pFAK described in TM cells and embryonic fibroblasts (51)(14). Consistent with biochemical analyses, TM cells showed I-V and pharmacological profiles characteristic of nonselective calcium-activated monovalent current (CAN) mediated by TRPM4 (i.e., inhibition by Ruthenium Red, 9-PA and shRNA). Its quasi-linearity points at the membrane presence of PIP2 (52) whereas inhibition by BAPTA indicates that the channel is activated within membrane microdomains within which TRPV4 and TRPM4 functionally interact. Ca^2+^ fluctuations and CAN were observed in ~50% of recorded cells, possibly because the cells were derived from multiple TM populations or existed in different stages of the cell cycle. An immortalized atrial cardiomyocyte cell line similarly showed TRPM4-dependent Ca^2+^ oscillations in ~30% cells (25).

The advantage of the synthetic agonist GSK1010 was that it allowed us to track signaling mechanisms associated with TRPV4 in the absence of concurrent activation of Piezo1 and TREK-1 SACs (e.g., (10)(11). In contrast to previous studies in myocytes, smooth muscle cells, neurons and immune cells (24)(25)(26)(53) which linked TRPM4-dependent [Ca^2+^]_i_ oscillations to obligatory voltage-gated Ca^2+^ influx, TM cells did not show functional VOCC expression under our experimental conditions. Thus, depolarization had no discernible effect on [Ca^2+^]_i_ and the transmembrane current, and Ba^2+^ concentrations that reliably trigger VOCC spiking and suppress store release in retinal neurons (36)(2) had no discernable effects on TM signaling. The absence of coupling between the membrane potential, [Na^+^]_i_ and Ca^2+^ oscillations indicates that Ca^2+^ fluctuations reflect intracellular and/or voltage-independent mechanisms. For example, previous investigation of TRPM4-TRPC3 interactions in HEK293 cells showed that cell depolarization caused by Na^+^ influx transiently suppresses TRP-mediated Ca^2+^ influx by reducing the inward electrochemical driving force for Ca^2+^ (43). In HL-1 cardiomyocytes, TRPM4 signaling, and oscillations were attributed to redistribution of mitochondrial Ca^2+^ stores into the cytosol (25) but it remains to be determined whether Ca^2+^ transients that underlie the oscillatory response involve *ψ_m_* and mitochondrial transporters (Na^+^/Ca^2+^-Li^+^ exchangers, Ca^2+^/H^+^ exchange, and/or MCU uniporters). Another potential source could include reciprocal interactions between Ca^2+^ influx and the IP3R store (54) mediated through contacts between the C-terminus CaM-binding site of TRPV4 and the IP3R receptor (55) as well as depolarization-dependent reduction of the driving force for Ca2+ influx through CRAC/Orai channels (24).

TRPM4 was suggested to be directly activated by membrane stretch (56) but this conclusion has been controversial (55)(57)(58). A compelling aspect of our findings is that its role in mechanosignaling might instead reflect activation downstream from SACs. The chemical activation of TRPV4 reproduced time-dependent behavior of pressure-induced Ca^2+^ events in human TM cells (44) and oscillatory phenotypes in TRPV4-expressing macrophages, fibroblasts, keratinocytes, endothelial cells, chondrocytes, and glia (46)(54)(46). Such oscillations have been linked to microcontractions, ECM remodeling and transfer of mechanical force across loadbearing focal adhesions (59)(45)(46). Given that TRPV4 and TRPM4 activation can augment expression of F-actin, alpha smooth muscle actin and fibrosis (60)(14), it is possible that stretch-sensitive tyrosine phosphorylations, RhoA signaling and contractility in TM cells (14) are dynamically regulated by TRPV4/TRPM4-dependent Ca^2+^ transients. Another intriguing possibility is that, as suggested for astroglial endfeet (54), TRPV4-dependent Ca^2+^ oscillations operate at the threshold of contractile (RhoA-dependent; (14) vs. relaxing (NO-dependent, (61) states that may be associated with myogenic-like contractility (62)(44)(63). Finally, TRPM4 shares the regulatory sulphonylurea receptor (SUR1/2) with inward rectifying K^+^ (Kir6.1 and Kir6.2) subunits that form K_ATP_ channels sensitive to [Ca^2+^]_i_ and intracellular ATP (64)(65). Efficacy of K_ATP_ channel openers in IOP lowering (66) might include their effects on the SUR1-TRPM4 complex, Ca^2+^ and Na^+^ homeostasis. We tested whether the oscillatory component associated with steady-state conventional outflow requires TRPM4 signaling but found a small, non-significant decrease in outflow facility. It remains to be seen whether the contribution of TRPM4 channels can be unmasked under conditions associated with maximal TRPV4 activation.

In conclusion, our findings provide support for the hypothesis that TM stretch sensing involves novel interactions between mechanotransducers, TRPM4 channels and Ca^2+^ stores that alter intracellular [Ca^2+^]_i_ and [Na^+^]_i_ gradients and can profoundly change the cells’ membrane potential. The persistent pacemaking loop between Ca^2+^ influx, release and Ca^2+^-activated channels activated by TRPV4 could, as by analogy with other load-bearing cells, be expected to time-dependently coordinate the activation of transcription factors, actomyosin-driven tension, mechanical load transfer across focal contacts, and secretion of matrix metalloproteinases. Although investigations of the TM typically focus on its smooth muscle cell- and endothelial-like properties, TM cells express immune-like markers such as complement C1QB, the tyrosine kinase binding protein (TYROBP), major histocompatibility complex proteins and the Toll-like receptor 4, which accord with their phagocytic, scavenger and immune functions. Future studies will show whether long-lasting TRPM4-dependent calcium oscillations during the calcium plateau phase, observed in TM cells, myocytes, acinar cells, mast cells and lymphocytes (24, 25,50), play comparable functions in phagocytosis, translayer Na^+^/water transport, and production of proinflammatory factors. Given that TRPV4 channels integrate a wide range of sensory inputs (mechanical stimuli, temperature, polyunsaturated fatty acids) known to modulate trabecular outflow, they provide a useful target for translational interventions aimed at finetuning IOP homeostasis.

## Acknowledgements

Supported by: NIH (R01EY027920, R01EY031817, P30EY014800, T32EY024234), Stauss-Rankin Foundation, USAMRAA and unrestricted support from Research to Prevent Blindness to the Moran Eye Institute at the University of Utah.

